# Sensing soluble uric acid by Naip1-Nlrp3 platform

**DOI:** 10.1101/2020.05.15.077644

**Authors:** Tarcio Teodoro Braga, Mariana Rodrigues Davanso, Davi Mendes, Tiago Antonio de Souza, Anderson Fernandes de Brito, Mario Costa Cruz, Meire Ioshie Hiyane, Dhemerson Souza de Lima, Vinicius Nunes, Juliana de Fátima Giarola, Denio Emanuel Pires Souto, Tomasz Próchnicki, Mario Lauterbach, Stellee Marcela Petris Biscaia, Rilton Alves de Freitas, Rui Curi, Alessandra Pontillo, Eicke Latz, Niels Olsen Saraiva Camara

## Abstract

The immune system can recognize microbes and sterile tissue damage. Among the damage-associated molecular patterns (DAMPs), uric acid is considered a major component which can trigger inflammation. It represents a breakpoint in the evolutionary history of humans as our ancestors lost the uricase gene, the enzyme responsible for its cleavage. High soluble uric acid (sUA) concentration is able to increase IL-1β in murine, but not human macrophages. We observed that sUA increased the mRNA expression of *Naip1* in murine macrophages, and, therefore, we hypothesized that the recognition of sUA can be made by a Naip1-Nlrp3 inflammasome platform. Additionally, we used genome-wide transcriptome analysis, functional analyses and structural modeling predictions and observed that virus-transduction of murine Naip1 into human macrophages induced IL-1β after sUA stimulus, besides leading to fatty acid production and an inflammation-related response. Moreover, pharmacologic inhibition and genetic loss of Nlrp3 led to decreased IL-1β production upon sUA stimulus. Surface plasmon resonance and quartz crystal microbalance showed that sUA is able to interact with Naip1. Naip could be a lost receptor for sUA in the evolutionary process and a better understanding of the immune modulatory function of sUA could lead to design rational novel anti-hyperuricemic therapies.

## Introduction

Host responses against harmful signals are basic physiological reactions of all living organisms. Innate immunity pattern-recognition receptors (PRRs) were firstly described as recognizing conserved structural components of microorganisms [1]. The discovery of Toll-like receptor (TLR) [2] led us to understand how the immune system responds to non-self-antigens in the context of an infection [3, 4], contrasting to the previous model in which the immune system reacted to all non-self-antigens while being tolerant to self-ones [5, 6]. Based on Polly Matzinger’s studies stating that “the immune system is more concerned with entities that do damage than with those that are foreign” [6], several damage-associated molecular pattern (DAMPs) have been described. Yet, according to this theory, the “foreignness” of a pathogen is not the important feature that triggers a response, and “self-ness” is no guarantee of tolerance. Indeed, receptors for endogenous and exogenous signals may have evolved simultaneously once vertebrates and pathogens have shared eons of evolutionary time and space [6]. Perhaps PRRs have not evolved to bind to pathogens at all; the pathogens, instead, may have evolved to attach to them and enhance their own survival [7], a hypothesis that would explain a puzzling feature of PRRs that each one can attach to many different kinds of molecules.

Among several DAMPs, uric acid (UA), the product of purine catabolism, released mainly from dying cells and ischemic tissues, is considered a major alarmin, especially when it is present at elevated levels and crystalized - also known as monosodium urate (MSU) [8]. In rodents, MSU activates the immune system [9, 10], acts as a pro-oxidant molecule, stimulates chemotaxis and also activates NF-κB and MAPK pathways [11]. Moreover, MSU induces the release of IL-1β through the activation of inflammasome-dependent caspases [10, 12, 13]. The inflammasome is a cytosolic complex mounted upon PAMPs/DAMPs sensing by a nucleotide binding domain (NBD/NACHT) and leucine-rich repeats (LRRs) containing receptor (NLR) [14]. NLRs belong to a superfamily of innate immune proteins with a very conserved structure along the phylogeny, from plants to mammals. NLRs shared NBD/NACHT and LRRs domains. Still they present a sub-family specific N-terminal domain: a pyrin domain (PYD) (NLRP), a caspase-recruitment domain (CARD) (NLRC), a Baculovirus Inhibitor of Apoptosis Protein Repeat (BIR) domain (NLRB), or an Acidic Transactivating Domain (ATD) (NLRA). In distinct phylogenic groups, the number of receptors and of paralogous genes differs, possibly as a consequence of a host/pathogen and/or environmental co-evolution [15]. Up to now, it has been demonstrated that both soluble UA (sUA) [16], as well as MSU, can induce Nlrp3 inflammasome activation in mice. In humans, gout is an inflammatory disease triggered by the deposition of MSU within joints and connective tissues, whereas Nlrp3 inflammasome is activated by UA crystals [17].

In humans, UA crystallization happens when its level reaches 6.8 mg/dL in plasma, while in rodents, the solubility threshold is about 10-fold lower [18]. Great apes have higher levels of UA in the serum (3.02 to 6.72 mg/dL, corresponding to 180 to 400 μM), compared to other animals (18 to 40 μM). This observation is compatible with the absence of the uricase (or urate oxidase) activity, the enzyme involved in purine catabolism converting UA into allantoin [19, 20]. The loss of uricase at the divergence between great apes and other mammals may be related to a survival advantage, as previously hypothesized, due to the UA characteristics as a molecule responsible for saving energy [21]; however, it takes up a tricky question about the role of a mammal’s sensor for UA. We hypothesize that along with uricase lost and a consequent elevation of UA serum levels; human has lost the sensors to recognize “high” levels of sUA.

Among NLR receptors able to induce inflammasome activation in mice and humans, the sub-family of NLRB took our attention. In mice, exist 6 paralogous genes, namely *Naip* 1-6 and 4 functional receptors (Naip-1, 2, 5, and 6), while in humans only one orthologous gene, *Naip* has been found [22]. Despite the fact that the differences in the aminoacidic content among the Naip proteins, both mice and human receptors have been described to play a role in defense against pathogens [23, 24]. Murine (m) Naip1 and Naip2 are respectively responsible for the detection of needle and rod proteins, structural proteins of the bacterial secretion system called type III secretion system (T3SS) [25–27]. Naip5 is responsible for the cytosolic recognition of flagellin [28], the major protein component of the bacterial flagellum. Similarly to Naip5, human Naip (hNaip) can bind bacterial flagellin [29] and to activate Nlrc4 inflammasome. In this scenario, Naip acts as a ligand sensor and the Nlrc4 is responsible for inflammasome activation and inflammasome-dependent cell death, known as pyroptosis [30]. Considering the differences of NLRB orthologous genes among different species and that until nowadays, any of NLRB was described as having endogenous ligands, in this study we performed transcriptome- and proteome-wide analysis in addition to interaction investigation and structural modeling predictions to study the sensing of sUA by mNaip1. Besides demonstrating that expressing mNaip1 into human cells allow them to be activated upon sUA stimulus, we found that mNaip1 directly recognize sUA. We then hypothesized that Naip could be the lost receptors for UA, and in particular for sUA.

## Results

### Naip1 is involved in sUA response

To assess the difference in serum basal level of UA among species, we have initially measured the UA levels of unrelated healthy adult human donors (n=5), C57Bl/6 adult mice (n=5), and adult old-world monkeys (rhesus macaque; n=5). As expected, humans presented an average blood UA concentration of 295 μM, 4, and 7 times more elevated UA levels when compared to mice and rhesus macaques, respectively (Fig. 1A). Then, we have stimulated murine LPS-primed BMDM and human and rhesus LPS-primed monocyte-derived macrophages with 200 μM of sUA. As observed, human cells did not produce IL1β after sUA stimulus when compared to LPS-primed cells (Fig. 1B). On the other hand, murine BMDM increased the IL1β production after sUA stimulus when compared to LPS-primed ones (Fig. 1C). Surprisingly, despite 200 μM being supra physiological level for rhesus macaque, their macrophages did not increase IL1β production after this sUA stimulus (Fig. 1D). Ischemic tissues consistently overproduce UA that trigger immune cell functions [31]. Both sUA stimulus and hypoxia condition of mouse BMDM led to increased Naip1 mRNA expression levels (5 and 15 times, respectively), but not Naip5 (Sup. Fig. 1A and 1B). Additionally, BMDM derived from Naip1^−/−^ and ΔNaip^−/−^ mice did not increase the IL-1β production upon sUA stimulation when compared to LPS-primed macrophages (Fig. 1E). On the other hand, Naip2^−/−^ and Naip5^−/−^ cells behaved as WT macrophages and, despite the variation within the group itself, there were no differences in the IL-1β production in the BMDM from Nlrc4^−/−^ animals after sUA stimulus in comparison with LPS-primed cells from Nlrc4^−/−^ mice (Fig. 1E). In an attempt to confirm the role of murine Naip (mNaip) platform into sUA response, we virus-transduced human THP1 cells with mNaip1, mNaip5, mNaip6, mNlrc4, and empty backbone vector. Our data demonstrate that PMA-activated and LPS-primed THP1 cells produced IL-1β after sUA stimulus only after mNaip1 transduction, but not mNaip5, mNaip6, mNlrc4 or the control empty vector (Fig. 1F). Such data point to mNaip1 as a target gene involved into sUA response.

**Figure 01.**
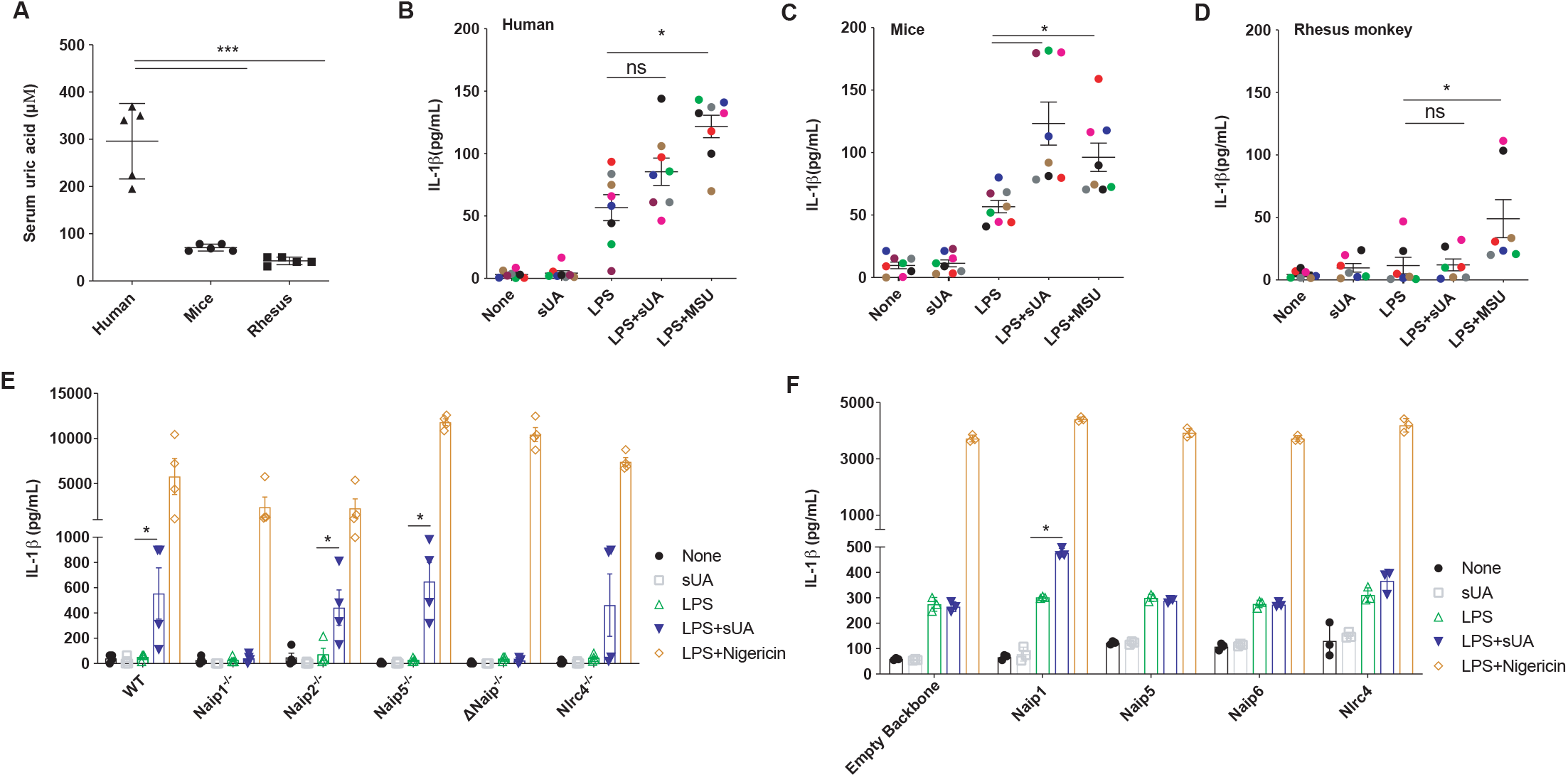
Human cells do not produce IL-1β upon sUA+LPS stimulus, unless they express mNaip1. (**A**) Uric acid levels measured in the serum of humans, mice and rhesus monkeys. IL-1β Elisa of (**B**) monocyte-derived macrophages collected from healthy people, (**C**) murine bone marrow-derive macrophages and (**D**) monocyte-derived macrophages collected from rhesus monkeys, all stimulated under different conditions. In B to D, LPS was added for 1 hour at 100 μg/mL and the media were posteriorly changed. MSU (100 μg/mL) and sUA (200 μM) were added for 6 hours. Each colored dot represents a different individual. (**E**) IL-1β Elisa of BMDM derived from Naip1^−/−^, Naip2^−/−^, Naip5^−/−^, ΔNaip^−/−^ and Nlrc4^−/−^ mice under different stimulus. (**F**) IL-1β Elisa of human THP1 cells virus-transduced with plasmids carrying Naip1, Naip5, Naip6, Nlrc4, and empty vector using lentivirus constructs and posteriorly stimulated under different conditions. In E and F, LPS was added for 1 hour at 100 μg/mL and the media were posteriorly changed. sUA (200 μM) were added for 6 hours and nigericin (10 μM) was added for 90 minutes. In A, n= 5 for each analyzed species. In B and C, we collected cells from 8 different individuals; in D, we collected cells from 7 different individuals. In E and F, data are plotted as median of a triplicate of three to four independent experiments. *p< 0.05, **p < 0.01, ***p < 0.001.

### IL-1β production in human cells is dependent on Nlrp3 and mNaip1

Naip1 carrying plasmid was modified by adding a “self-cleaving” 2A (T2A) sequence between Naip1 and the color-tag sequences. In this sense, the overexpressed Naip1 protein is not color-tag, which could implicate in some misinterpreted data. Initially, it was investigated the ability of PMA-activated, LPS-primed, and mNaip1-expressing THP1 cells to produce IL-1β under the stimulus of UA degradation products, i.e. allantoin, urea and ammonium. None of the investigated products were able to induce IL-1β production, as sUA did (Fig 2A). The role of Nlrp3 for sUA sensing, as previously stated [16], was further investigated. For that, we initially evaluated the IL-1β production upon sUA stimulus in THP1 cells virally transduced with either mNaip1 or empty backbone, both stimulated in the presence or absence of a Nlrp3 inhibitor, CRID3 (1 μM) [32]. IL-1β levels are reduced in cells pre-treated with CRID3 (Fig. 2B). IL-1β production was also evaluated in THP1 cells after Nlrp3 gene deletion by Crispr-Cas9. It was confirmed that IL-1β production is dependent on Nlrp3 activation (Fig. 2B-D) once Nlrp3-deleted cells exhibited decreased levels of IL-1β upon sUA stimulus (Fig. 2C and 2D). To investigate the interaction between Naip1 and Nlrp3, an immunoprecipitation assays with THP1 cell lysates using both Nlrp3 and Naip1 as targets was performed (Sup. Fig. 2). However, the targets were only found in the whole-cell lysates but not in the immunoprecipitants, suggesting that Nlrp3 and Naip1may not directly interact. Altogether, these data indicate that the observed IL-1β production followed by sUA stimulus requires both Naip1 and Nlrp3 inflammasome platforms.

**Figure 02.**
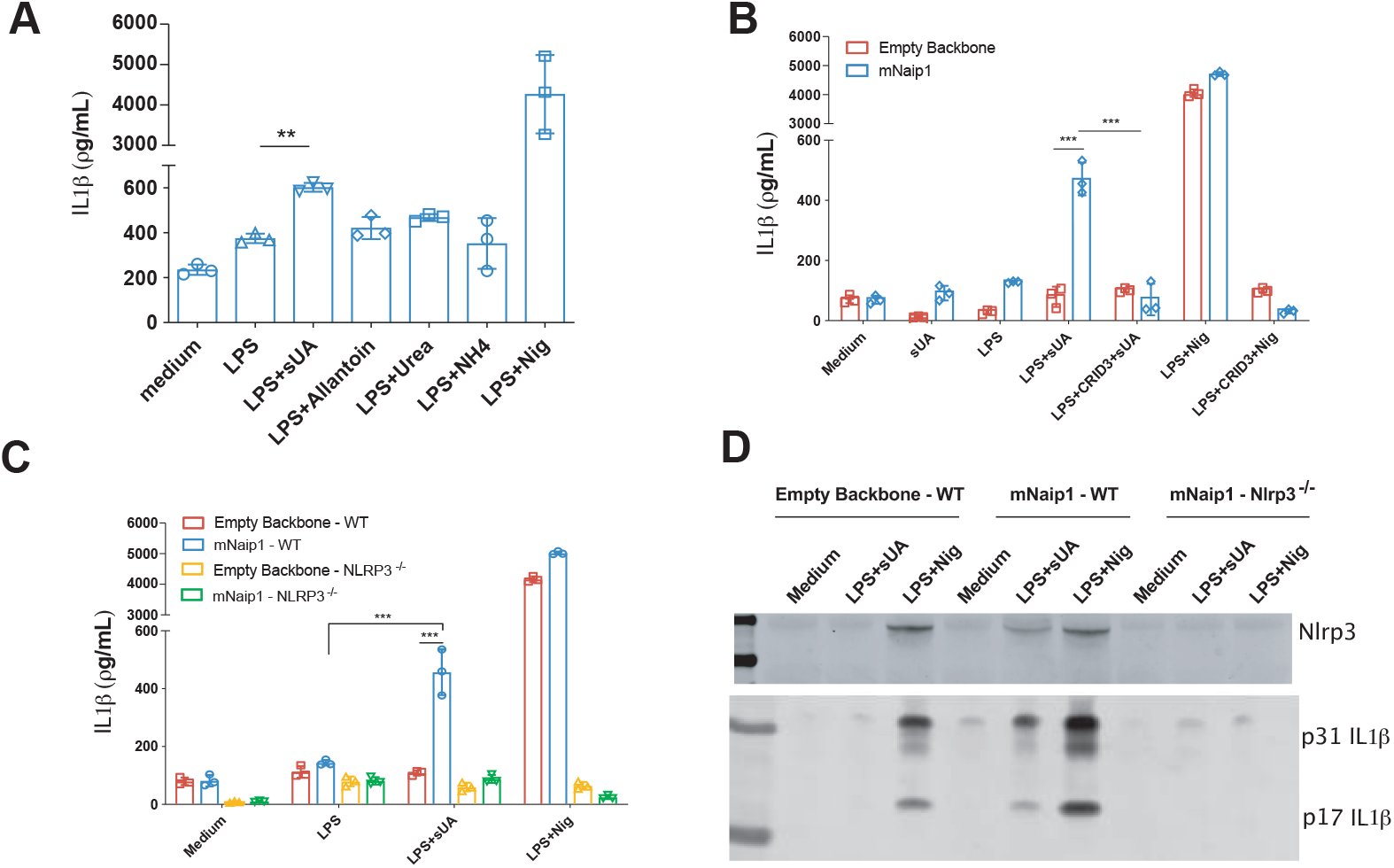
Naip1 and NLRP3 are required for LPS-primed THP-1 cells to produce IL-1β upon sUA. (**A**) IL-1β Elisa of mNaip1 transduced LPS-primed THP1 cells, stimulated with the products of uric acid degradation allantoin, urea and ammonium, the control non-treated cells (Medium) and LPS-primed treated with nigericin. (**B**) IL-1β Elisa of THP1 cells virus-transduced with empty backbone or with mNaip1 after 1 hour pre-treatment with LPS (1 μg/mL), 6 hours treatment with sUA (200 μM), 30 minutes treatment with nigericin or control non-treated cells (Medium). Some groups were pre-treated with the Nlrp3 inhibitor CRID3 at 1 μM 30 minutes before LPS priming. (**C**) IL-1β Elisa of WT THP1 and Nlrp3^−/−^ THP1 cells virus-transduced with empty backbone or with mNaip1 after 1 hour pre-treatment with LPS, 6 hours treatment with sUA, 30 minutes treatment with nigericin or control non-treated cells (Medium). (**D**) IL-1β and Nlrp3 western blotting images of WT THP1 and Nlrp3^−/−^ THP1 cells virus-transduced with empty backbone or with mNaip1 in control non-treated condition (Medium), or LPS-primed and treated with sUA for 6 hours, or with nigericin for 30 minutes. In D, data are representative of three independent experiments. All experiments were performed three different times and data are plotted as median of a triplicate. *p < 0.05, **p < 0.01, ***p < 0.001.

### Naip1 triggers enhanced immune responses and altered cellular metabolites content toward sUA

To better define how mNaip1 influences sUA sensing, RNA-seq analysis of mNaip1- and backbone-transduced THP1 cells was performed. The presence of Naip1 affects the gene transcription, which was evident when the gene expression differences are represented as a Volcano plot (Sup. Fig. 3A). Unsupervised hierarchical clustering of 8,000 genes (Sup. Table 1) and the top 20 most down and up-regulated genes by means of centered logarithm of FPKM values in the six replicates of each experimental group demonstrated that sUA-stimulated macrophages globally reprogram transcriptional responses after expressing mNaip1 (Sup. Fig 3B). Detailed inspection of the most highly differentially expressed genes revealed that *ccl2*, *pik3cd*, *nck2*, *tab1*, and *fgfr1* were expressed more strongly in mNaip1-expressing GFP-tagged cells stimulated with sUA, when compared to GFP-transduced (control) cells at the same stimulus (Sup. Fig. 3C). Functional annotation enrichment analysis for KEGG and PANTHER pathways demonstrated that PMA-activated and LPS-primed THP1 cells expressing mNaip1 lead to a shift toward increased inflammation, cancer- and infection-related signaling pathways (Sup. Fig. 3D). Enrichment term analysis using the genes that were upregulated in mNaip1-expressing macrophages, was further visualized as a KEGG pathway enrichment network (Sup. Fig. 3E). Together, these analyses identified that upregulated genes were associated with processes involved in cytoskeleton regulation, adherent junctions, proteoglycans in cancer, as well as bacterial infection and invasion, and immune processes (Sup. Fig. 3E). These data demonstrate that mNaip1 triggers enhanced immune responses after sensing sUA.

In addition to gene expression profile, we performed a proteomic analysis to identify differentially expressed proteins followed by sUA stimulus in both control (GFP-transduced) and mNaip1-expressing LPS-primed THP1 cells, and we highlighted in orange the proteins only found upon sUA stimulus and in yellow the proteins found only in LPS-primed condition (Sup. Fig. 4A, Sup. Table 02 and Sup. Table 03). Moreover, the two different cells under sUA stimulus were compared (Sup. Fig. 4B). Among 44 proteins only present in mNaip1-expressing cells, but not present in control ones, we highlighted 30 proteins, including some related to immune response such as thymopoietin (A0A024RBH7), CD99 (A8MQT7), stress-associated endoplasmic reticulum protein (Q9Y6X1), and lysosome-associated membrane glycoprotein 2 (H0YCG2) (Sup. Fig 4B). Additionally, it was also observed an up-regulation of mitochondrial citrate synthase (F8VRP1), amino acid transporter (M0R106), mitochondrial glutamate carrier 1 (Q9H936), inorganic pyrophosphatase (Q15181), alpha-N-acetylgalactosaminidase (P17050), phosphoinositide phospholipase C (B7Z5V4), and acyl-CoA dehydrogenase, isoform 4 (A0A0S2Z3A5), all associated with cellular metabolism (Sup. Fig 4B). On the other hand, we highlighted 60 proteins among 342 ones only present in GFP^+^ control cells, such as transforming growth factor-beta-induced protein (H0Y8L3), solute carrier family 25-member 4-isoform 3 (A0A0S2Z359), vimentin (B0YJC4), among others (Sup. Fig 4B and Sup. Table 04). Of note, 84% of the analyzed proteins in the comparison between the two different cells under sUA stimulus were detected in the RNASeq data.

However, only 2.8% of the expressed proteins correspond to differentially expressed genes (data not shown). We built a STRING network view of proteins only present in mNaip1 expressing cells when compared to GFP^+^ control ones, both under sUA stimulation (Sup. Fig. 4C), which evidenced the metabolism-related up-regulation pathways, especially those related to lipid metabolism. These data demonstrate that additionally to immune response, mNaip1 triggers altered cellular metabolites content in LPS-primed macrophages stimulated with sUA.

### Naip1 activation may be potentiated after the elevation of the cellular content of total lipid

Following the altered cellular metabolites content, UA is also described as increasing the accumulation of triglyceride into hepatic cells [33]. In this sense, we next investigated lipid drops formation in LPS-primed THP1 cells stimulated with sUA. It was observed an increase in cellular content of lipids after sUA stimulus but in a mNaip1-independent way (Fig. 3A-B). Alteration in metabolites content could also change mitochondrial activity once these plastic organelles sense cellular metabolites, oxygen, and nutrients, and they exert central roles as source of energy and ROS [34]. Changes in mitochondrial membrane potential in live cells upon sUA stimulus were therefore measured (Fig. 3C-D). Despite no differences in mitochondrial area within mitotracker staining, a reduced mitochondrial membrane potential was observed, as indicated by failure to load the positively charged mitochondrial indicator TMRE in mNaip1 expressing cells stimulated with sUA, when compared to LPS-primed cells (Fig. 3C-D). We next measured the oxygen consumption ratio (OCR) of THP1 cells virally transduced with empty backbone or mNaip1, both LPS-primed, treated or not with sUA. sUA increased OCR but in a Naip1-independent manner. The mitochondrial pyruvate carrier inhibitor UK5099 (100 μM) was used in order to evaluate the ATP consumption derived from fatty acid indirectly. Again, such OCR increasing can be reverted by UK5099 pre-treatment in both cell types (Sup. Fig. 5A and 5B). Altogether, such data indicate that mNaip1 expression alters the cellular fatty acid content and reduces active mitochondria number upon sUA stimulus.

**Figure 03.**
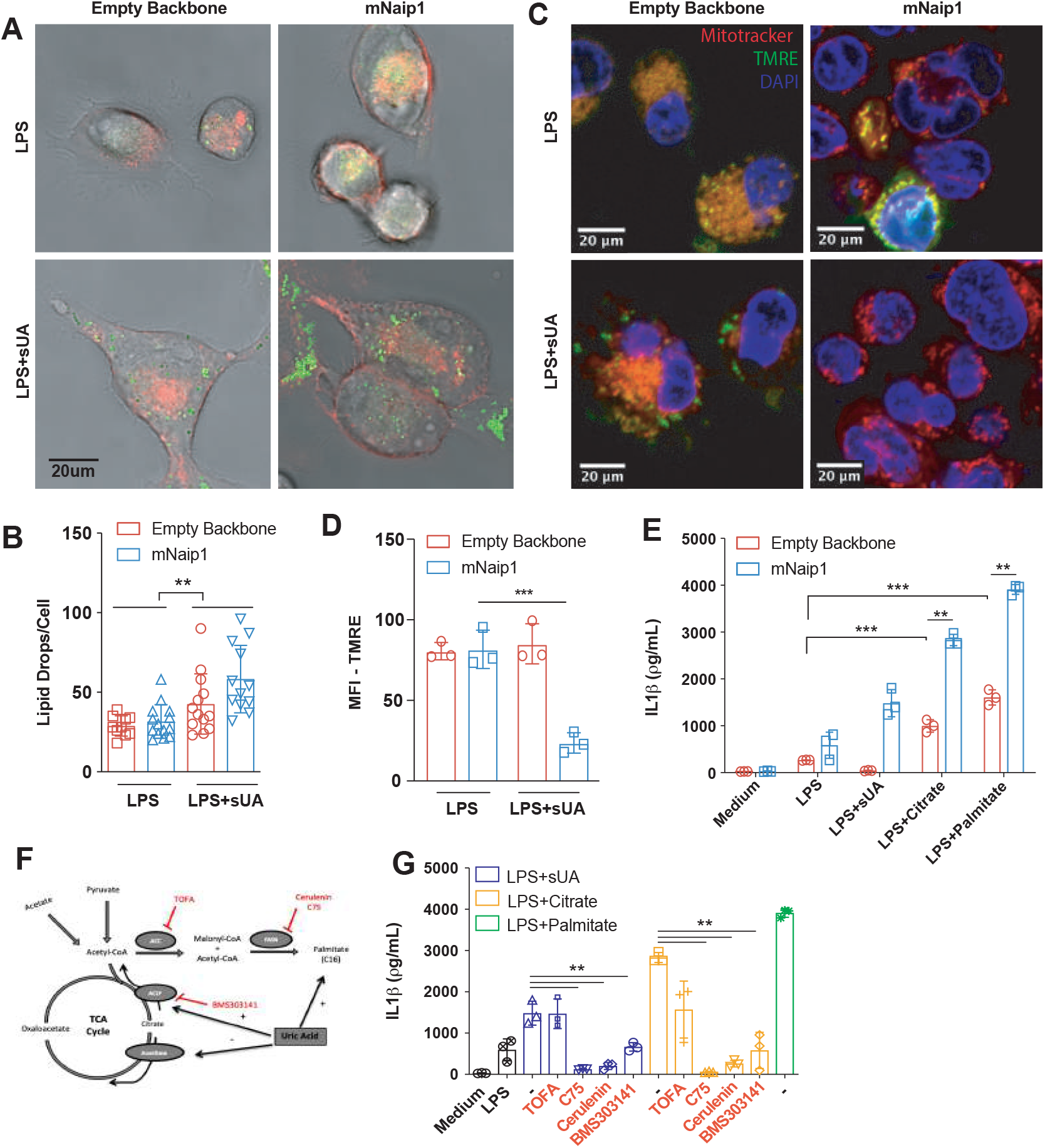
Naip1 activation upon sUA stimulus may be potentiated after elevation of the cellular content of neutral lipid. (**A**) Representative images of THP1 cells transduced with empty backbone or mNaip1 at LPS-primed condition or LPS-primed and stimulated with sUA for 6 hours. The membrane is in red and lipid droplets are stained for LD540, in green. (**B**) Quantification of lipid droplets per cell. (**C**) Representative images of empty backbone- and mNaip1-transduced cells primed with LPS or LPS-primed and sUA-stimulated stained with mitotracker (red), tetramethylrhodamine ethyl ester (TMRE) (green) and DAPI (blue). The bars in each image represent 20 μm (**D**) TMRE quantification, indicating polarized mitochondria of the experiments in C. (**E**) IL-1β Elisa of empty backbone- (red bars) and mNaip1- (blue bars) transduced cells, both at non-stimulated (Medium) condition or LPS-primed and stimulated for 6 hours with sUA, citrate or palmitate. (**F**) Schematic representation of the TCA cycle and the fatty acid synthesis pathway given emphasis to the inhibitors and stimulus used in J. (**G**) IL-1β Elisa of mNaip1 transduced and LPS-primed cells, stimulated for 6 hour with sUA (200 μM), citrate (5 mM) or palmitate (100 μM), in the presence or absence of ATP citrate lyase inhibitor (BMS303141 at 25 μM), acetyl-CoA carboxylase inhibitor (TOFA at 10 μg/mL), or fatty acid synthase inhibitors (C75 at 50 μM or Cerulenin at 5 μg/mL). In A and B, data are representative of three independent experiments and n = 12. In D, data are plotted as median of ten different micrography fields of three independent experiments. In E and F, the experiments were performed three different times and n=3. *p < 0.05, ***p < 0.001.

We next investigated whether the elevation of fatty acid synthesis could trigger IL-1β production. For that, it was measured the levels of IL-1β in the supernatant of LPS-primed cells virally transduced with empty backbone or mNaip1 after 6 hours of incubation with citrate [35] and palmitate. Despite significant production of IL-1β into empty backbone-transduced cells upon both citrate (5 mM) or palmitate (100 μM) stimulus when compared to LPS-primed cells, mNaip1-expressing cells produce even higher levels of IL-1β when compared to empty backbone transduced cells regardless the stimuli (Fig. 3E). Moreover, LPS-primed mNaip1-expressing cells were stimulated with sUA or citrate in the presence of the acetyl-CoA carboxylase-α inhibitor, TOFA (10 μg/mL), or the phosphor-ACLy inhibitor, BMS303141 [36] (25 μM), or the fatty acid synthase inhibitors, C75 (50 μM) and cerulenin (5 μg/mL) [37], as shown in the schematic figure 3F. As observed, all these inhibitors, but TOFA, led to decreased levels of produced IL-1β (Fig. 3G) upon sUA or citrate stimuli. Altogether, our data suggest that sUA leads to fatty acid synthesis in a way independent of mNaip1. Saturated fatty acids promote Nlrp3 inflammasome activation [38], especially, palmitate [39]. We added that citrate- and palmitate-mediated IL-1β production is potentiated in the presence of Naip1.

### Naip1 directly recognizes sUA

In a study investigating the role of different lipids in macrophage lipidomic, palmitate presented the most pronounced effects [40]. In order to investigate whether mNaip1 directly senses sUA and/or palmitate, we performed a quartz crystal microbalance with dissipation (QCM-D) analysis. After an initial immobilizing step with anti-GFP, we incubated sUA solution sample in crescent concentrations (12.5 μM; 25 μM; 50 μM; 100 μM; and 200 μM). It was observed that mNaip1 directly interacts with sUA. As sUA adsorption was reached, the QCM-D frequency changed at the maximum, even at the lowest studied concentration (Fig. 4A), indicating saturation at lower concentrations, and an increase in the mass adsorbed per area. Yet, we compared the interaction of mNaip1 with sUA and with palmitate in real time in a surface plasmon resonance (SPR) immunosensor. With SPR-based biosensor technology, mNaip1 is tethered to the surface of a previously functionalized SPR sensor chip and the possible ligands are introduced in solution, as illustrated in the scheme of figure 4B. The SPR curve (sensorgram) obtained in real time for all steps involved in the evaluation of interactions between sUA and mNaip1 protein (purple curve), and between palmitate and mNaip1 protein (green curve) is observed in figure 4B. It is important to observe a significant variation of response (Δθ_SPR_) obtained after addition of sUA (2 μM), which characterizes the interaction between sUA and mNaip1 protein. In this phase of the study, higher concentrations of sUA (12.5 to 200 μM) were also accompanied by significant responses (data not shown). In turn, the addition of palmitate at the concentration of 2 μM (green curve) and at higher concentrations (12.5 to 200 μM-data not shown) did not trigger a notable response. These results suggest that sUA directly binds to the mNaip1 protein.

**Figure 04.**
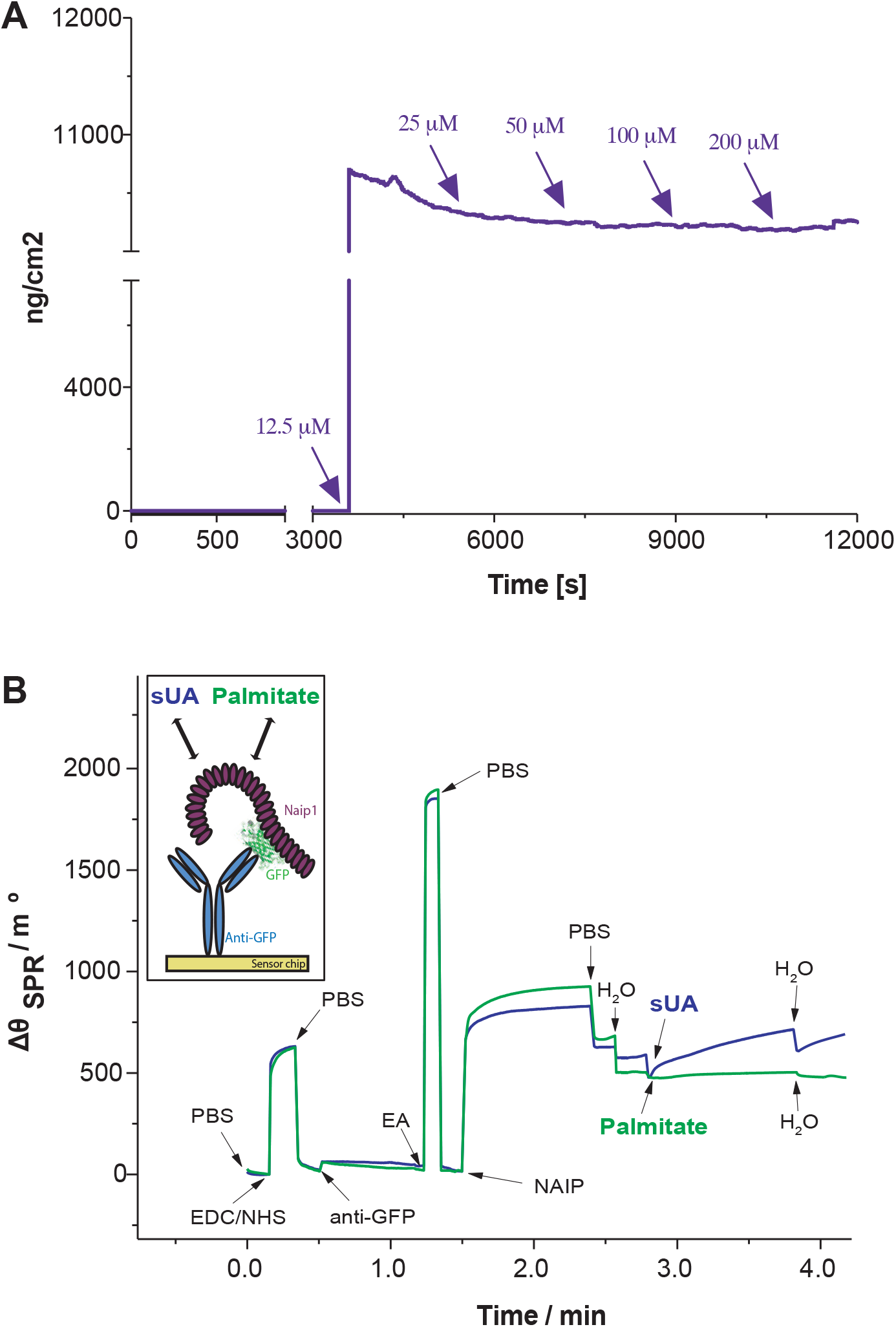
QCM monitoring and SPR sensorgram evidencing all steps involved for the detection of the interaction between sUA and Naip1 protein. **(A)** QCM responses over time within sUA injection after Naip1 immobilization upon anti-GFP adsorption on the gold quartz crystals surface at 37°C. The arrows indicate sample injection. **(B)** Schematic representation of the constructed SPR sensor chip (in the box). Sequential addition of compounds into the system: (i) addition of the buffer solution (PBS, 10 mmol L^−1^ at pH 7.4); (ii) mixture consisting of EDC (150 mmol L^−1^) and NHS (150 mmol L^−1^); (iii) PBS; (iv) immobilization of anti-GFP (10 μg mL^−1^); (v) PBS; (vi) addition of ethanolamine (EA); addition of cell lysates containing Naip1 protein (2 μg mol L^−1^). It is possible observe a very intensive response for the interaction of the Naip1 protein with anti-GFP; (v) PBS;(vi) addition of pure H_2_O; (vii) addition of sUA (2 μmol L^−1^, purple line) and palmitate (2 μmol L^−1^, green line). It is possible to observe the significant variation of the SPR angle (Δθ_SPR_) due to the interaction between sUA and Naip protein. In A and B, data are representative of three independent experiments.

### hNaip and mNaip1 may respond differently to uric acid

As previously studied, for inflammasomes to be formed, NLR proteins, like Naip, must recognize ligands to be released from their autoinhibited state to finally trigger the oligomerization of NLRCs, and assemble inflammasome complexes [41, 42]. After modelling, the structures of mNaip1 and hNaip, in their inactive forms, we observed important differences on their surface electrostatic properties, which may directly interfere with their ability to recognize specific ligands. By performing molecular docking, we investigated regions of possible binding of sUA onto the solvent accessible surface of both Naip proteins (Fig. 5). More than 250 iterations were performed for each target. Given the ligand binding geometry in each interaction, the docking results were summarized as clusters containing one or more ligands with similar bind poses. The lower the Gibbs free energy (ΔG) of a ligand-target interaction, the higher is the affinity between the interactors, and for comparison purposes, figure 5 only displays clusters with ΔG lower than −6 kcal/mol, with the binding pose with lowest energy (i.e. highest affinity/stability) being highlighted in both targets. As expected due to their structural differences, the regions where sUA binds in both Naip proteins mostly disagree. While the predicted binding region of sUA in mNaip1 lies on the NBD domain (ΔG = −7.92 kcal/mol) (Fig. 5A), for hNaip the ligand most likely exhibits higher affinity to the LRR domain (ΔG = −8.00 kcal/mol) (Fig. 5D).

**Figure 05.**
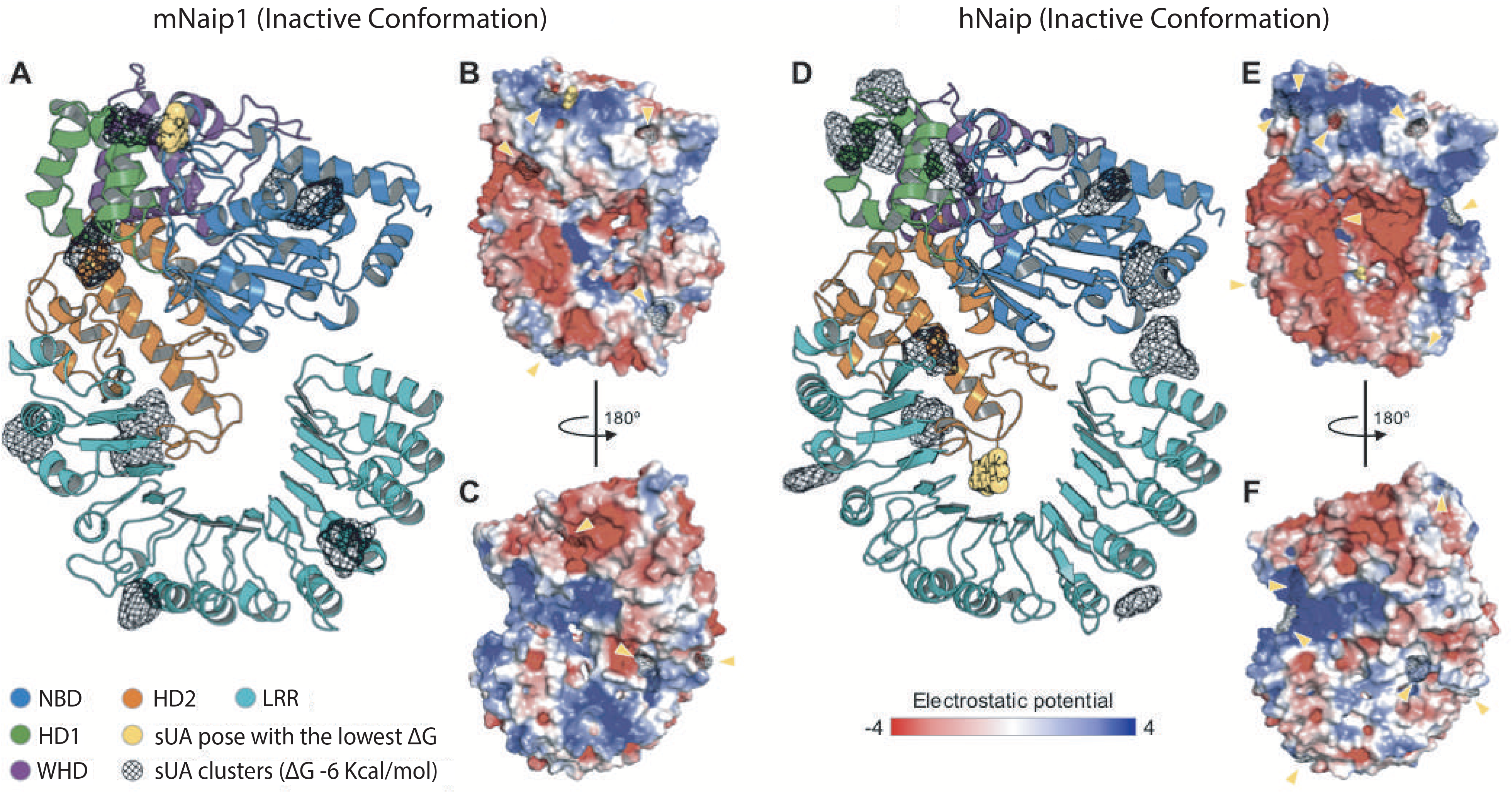
Structural analysis of the inactive conformations of hNaip and mNaip1, modelled by homology using an inactive form of Nlrc4 structure (PDB 4KXF) as template. **(A)** Cartoon representation of a mNaip1 (Uniprot: Q9QWK5) homology model. The distinct colours represent functional regions commonly found in proteins of the NLR family (NBD-HD1-WHD-HD2-LRR), coloured as shown in ZHANG, 2015. Clusters of uric acid (URC) molecules are shown as black meshes, which represent points on the Naip surface where two or more URC were found to bind, in multiple independent rigid docking simulations. The pose with the lowest ΔG is shown as a yellow sphere representation. **(B)** Surface electrostatic potential calculated for mNaip1. Its solvent accessible surface is shown with a potential gradient ranging from < −4 kBT (red) to > 4 kBT (blue). Yellow arrows highlight URC clusters shown in (A). **(C)** 180-degree rotation of mNaip1 around its Y axis. **(D)** Cartoon representation of a hNaip (Uniprot: Q13075) homology model. **(E**) Its surface electrostatic potential. **(F)** 180-degree rotation of hNaip around its Y axis. See legend for more details.

In association with our experimental results, based on homology modelling and molecular docking, we hypothesize the surface electrostatics of mNaip1 enables this protein to recognize sUA and stop its autoinhibition, tasks that hNaip is unable to perform. With the deletion of the uricase gene on great apes [19], over its evolution, mutations on hNaip surface electrostatics were probably selected to increase its physiological tolerance to high levels of sUA, in such a way to prevent activation of inflammasomes, and allow great apes to benefit from the survival advantages provided by high levels of serum UA [21].

## Discussion

Evidence is reported that sUA increases Naip1 transcription in murine macrophages. So, Naip1 could be responsible for cellular signalling triggered by sUA. Among all mNaip, Naip1 presents the higher aminoacidic content similarity with the hNaip, about 70%. Despite some reports pointing to hNaip recognizing the same ligand as mNaip5 [29], some studies indicate that hNaip recognizes the same bacterial components as mNaip1 [26, 43]. It is possible that the 30% differences between human and murine proteins are associated with sUA signaling. This is the first study to postulate that Naip recognizes a DAMP. Once our previous work suggests that sUA activates the Nlrp3 inflammasome [16], we also investigated the role of Nlrp3 in mNaip1 expressing cells. LPS-primed human THP1 cells only produce the mature form of IL-1β under sUA stimulus when they express Nlrp3 and mNaip1. Most inflammasomes are believed to include only a single NLR, though other NLR-NLR interactions have been proposed. The interplay between Nlrp3 and Nlrc4 reveals an unexpected overlap between what had been considered distinct inflammasome scaffolds [44]. Besides, it was reported that Nlrc4 can recruit Nlrp3 through its NACHT domain, in the context of *S. typhimurium* infection [44]. It was also demonstrated that NEK7, a member of the family of mammalian NIMA-related kinases (NEK proteins), is a Nlrp3-binding protein that acts downstream of potassium efflux to regulate Nlrp3 oligomerization and activation [45]. Hence, besides the mechanisms by which sUA activates mNaip1 inflammasome, it remains to be determined if mNaip1 interacts with Nlrp3 after sUA stimulus once we observed no clear evidence of Nlrp3 and mNaip1 protein interaction.

Besides responding to PAMPs and DAMPs, recent data suggest that the immune system act as a signal integrator able to detect disturbances in cytoplasmic cells related to metabolites. This monitored disruption is termed "homeostasis-altering molecular processes" (HAMPs) [46], and it provides powerful flexibility in the ability of the innate immune system to detect infections and chronic inflammatory diseases. It has been shown that the Nlrp3 inflammasome complex activation and the posterior caspase-1 and IL-1β production occurs following the saturated fatty acid palmitate triggering even in humans’ cells [47, 48]. Both QCM and SPR immunosensor based on the immobilization of Naip1^GFP^/anti-GFP/Au for specific sUA detection were successfully performed. The results indicate that the three-dimensional structure of mNaip1 provided a great accessible area for interaction with sUA, and no accessible area for interaction with palmitate. Homology modelling and molecular docking analysis indicate that the surface electrostatics of mNaip1 enables it to recognize sUA through abolishing its autoinhibition state, tasks that hNaip is unable to perform. Despite we have demonstrated that sUA, but not palmitate, is responsible by directly binding to mNaip1, our data pointed to an increased IL-1β production in the context of Naip1 expression in human’s cells followed by palmitate stimulus. Therefore, further analyses are necessary in order to better investigate the mechanisms by which palmitate could lead to Naip1 permissiveness to sUA.

Additionally, the transcriptome and the cellular metabolites content was changed in cells upon sUA stimulus, being the presence of mNaip1 altered some metabolism-related enzymes besides favoured the increase in the immune responses toward sUA. Based on the central dogma, it was generally assumed that there exists a direct correlation between mRNA transcripts and protein expressions, however, recent studies have been showing that this correlation can be low due to various factors such as post transcription modifications [49] or even the time mRNA and protein are expressed [50]. Moreover, we observed it reduced active mitochondria in mNaip1 expressing cells stimulated with sUA. This result corroborates a study demonstrating that the percentage of TMRE^+^ cells was significantly lower in LPS-primed macrophages stimulated with ATP, compared to the control ones [51]. In this mentioned study, the TMRE dye was not trapped in the mitochondrial membrane due to depolarization consequent to calcium release.

Uricase activity missing through the evolutionary process gave UA a puzzling character in the evolutionary history of Humans. Great apes have, in the basal state, high physiological levels of sUA and humans’ macrophages do not respond to 200 μM sUA, an inflammatory condition for murine cells. Uricase inhibition therapy and the consequent elevation in the serum UA is responsible for the triggering metabolic syndrome comorbidities in murine models [52–54]. We have demonstrated, on the other hand, that rhesus macaque, a primate with uricase enzyme activity evolutionarily maintained, has reduced levels of IL-1β production by their monocyte-derived macrophage following sUA stimulation. It is possible that rhesus’ macrophages require a higher priming activation in order to induce IL-1β transcription since LPS alone could not increase IL-1β production in these experiments. Moreover, several cytokines, including IL-1β, circulate at very low levels in both affected and unaffected rhesus macaques, in a different model of diseases [55–57]. It is also speculated that rhesus Naip protein was selected to tolerate elevated levels of serum UA.

In recent years, an understanding of additional adverse effects of high levels of serum UA has been advanced [58]. Early scientific literature suggested an association between uric acid concentration and incidence of cardiovascular disease, specifically, the development of hypertension [59], metabolic syndrome [60], endothelial dysfunction [61], and microalbuminuria [62]. Lifestyle and socioeconomic changes that occurred over time have resulted in a marked reduction of physical activity as well as in profound dietary changes. These changes correlate to increased rates of metabolic diseases triggered by overly active innate immune functions [63], being the chronic inflammation termed ‘metaflammation’ [64, 65]. Furthermore, multiple genetic and non-genetic risk and protective factors are also thought to contribute to the pathogenesis of metabolic diseases, specifically those related to hyperuricemic conditions. Different states of tolerance to sUA sensing by hNaip could predict innate immune activation state in the context of hyperuricemic-related diseases. In this sense, besides understanding the humans evolutionary process, investigating which mechanisms mediate the immune modulatory function of sUA is also essential to better design rational novel anti-inflammatory therapies.

## Materials and Methods

### Soluble uric acid preparation

Media was pre-warmed (37°C), uric acid (Ultrapure, Sigma; 200 μM) was added and then sterilized through 0.20 μm filters. Crystals were not detectable under these conditions (polarizing microscopy), nor did they develop during cell incubation.

### Reagents

Ultrapure LPS was obtained from InvivoGen, nigericin was obtained from Invitrogen. DRAQ5 was purchased from eBioscience. Anti-GFP monoclonal antibody (GF28R) was from ThermoFisher Scientific. CRID3 was from R&D Systems, Allantoin, Urea and Palmitate were from Sigma-Aldrich. TOFA (5-(tetradecyloxy)-2-furoic acid) (CAS 54857-86-2) was purchased from Abcam. C75 (CAS 191282-48-1) was also from Abcam. Cerulenin (17397-89-6) was from Sigma-Aldrich, and BMS303141 (CAS 943962-47-8) was from Cayman Chemical. UK5099 CAS 56396-35-1 was from Merck Millipore. Ethanolamine (EA), 11-mercaptoundecanoic acid (11-MUA), N-(3 dimethylamino-propyl)-N-ethylcarbodiimidehydrochloride (EDC), and N-hydroxysuccinimide (NHS), were obtained from Sigma-Aldrich Chemical (St. Louis, MO, USA).

### Macrophages obtainment

Primary murine macrophages were generated from bone marrow. Briefly, bone marrow-derived cells were filtered through sterile polystyrene syringes (70–100 μm) and divided into 10 Petri dishes (100 × 20 mm; BD, Franklin Lakes, USA) supplemented with murine M-CSF (20 ηg/mL; R&D Systems, Minneapolis, MN, USA) diluted in DMEM-High (Invitrogen, USA) in the presence of 5% fetal bovine serum for 7 days. All procedures with mice were approved by the local ethics committees at the University of São Paulo (Document 45/2009). All experiments were performed in accordance with relevant guidelines and regulations.

Human and rhesus macaque (*Macaca mulatta*) macrophages were generated from monocyte-circulating cells. A centrifugation gradient obtained human and monkey peripheral blood mononuclear cells (PBMCs) in Ficoll-Paque (Dominique Dutscher). Subsequently, PBMCs were centrifuged in 51% PBS-diluted Percoll (GE Healthcare Life Sciences), and monocytes (> 70% purity) were adhered in six-well plates for 4 hours. After removal of non-adherent cells by washing the plates, monocytes were differentiated into macrophages in RPMI medium (Gibco, Grand Island, NY, USA) supplemented with 10% FCS (Gibco) plus antibiotics and antimycotics (100 U/mL penicillin, 100 g/mL streptomycin, and 25 g/mL amphotericin; Gibco) in the presence of human M-CSF (25 ηg/mL; R&D Systems, Minneapolis, MN, USA) for five days. All procedures with rhesus macaque were approved by the Ethic Committee on Animal Use of the Butantan Institute (CEUAIB) under the number 9376040717. Leucocytes’ concentrates were obtained after plasmapheresis at the Blood Bank Service of the “Hospital das Clinicas” in Sao Paulo (SP, Brazil) and they were used for peripheral blood monocytes isolation. All volunteers signed informed consent in compliance with the respective Institutional Ethics Committee.

### Viral transduction

Plasmids for GFP-tagged murine Naip1 (#60200), Naip5 (#60205), Naip6 (#60202), Nlrc4 (#60199) and empty vector (#60206) were purchased from Addgene (Watertown, MA, USA). Transformed bacteria with GFP-tagged murine Naip2 (#60201) plasmid were not able to grow. Plasmid #60200 was additionally modified either by fusing Naip1 with GFP or by adding a “self-cleaving” 2A (T2A) sequence between Naip1 and the color-tag sequence, with standard cloning techniques. Plasmid #60206 was used as control empty backbone. Briefly, THP-1 cells (ATCC, TIB202) were retrovirally transduced with constructs for the indicated plasmids. After retroviral transduction, cells were flowed cytometrically sorted to similar levels of GFP expression. Nlrp3-deficient (Nlrp3^−/−)^ macrophages have been previously described [66], and the Nlrp3 deletion into THP-1 deletion was performed according to Schmid-Burgk and colleagues [67].

### Cytokine profile

Cells lysates were maintained at RIPA buffer with protease inhibitors, at −80°C until dosage. IL-1β protein was measured using IL-1β (R&D Systems, Minneapolis, MN, USA), according to the manufacturer’s instructions.

### RNA extraction, library construction and sequencing

Total RNA was extracted from GFP-sorted THP1 cells containing lentiviral vectors NAIP1 (n=6), or empty vector control GFP^+^ cells (n=6) using TRIzol reagent according to manufacturer instructions (Thermo Fisher Scientific, Waltham, MA, USA). Integrity of total RNA was checked using Bioanalyzer 2100 (Agilent Technologies, CA, USA) with RNA Nano 6000 kit. Purity and quantity were measured using NanoDrop 1000 spectrophotometer and Qubit RNA HS fluorescence kit (Thermo Fisher), respectively. Total RNA concentration of each sample (n=12) was adjusted to 1 μg and used for poly (A) mRNA enrichment and stranded specific RNA-Seq library preparation with TruSeq Stranded mRNA kit (Illumina, San Diego, CA, USA). Quality control of prepared libraries were performed using Agilent Bioanalyzer DNA 1000 kit and Qubit DNA HS fluorescence kit. Libraries were pooled then sequenced at CEFAP-USP (Sao Paulo, Brazil) on an Illumina NextSeq platform on 75-bp paired-end mode using a High Output Kit.

### RNA-Seq data analysis

Before read mapping, clean reads were selected after preprocessing with Trimmomatic [68] removing adapter and poly-N sequences. After cleaning, the quality of reads was checked by FastQC tool then aligned to the human genome (GRCh38/hg38) using HISAT2 aligner (V2-2.0.0) [69] considering strandness. Overall mapping quality and uniformity of read coverage on exons were checked by RSeQC tool to ensure good RNA integrity and reproducible RNA sequencing. Stringtie (v.1.3.4) [70] and Ballgown [71] algorithms were applied to identify significantly differentially expressed genes (q-value < 0.05), based on the “new Tuxedo” package [72]. Gene set enrichment analyses were performed using Enrichr tool and GAGE package using up-regulated (FC > 1 and q-value < 0.05) and down-regulated genes (FC < 1 and q-value < 0.05).

### Proteomics analysis

Protein solutions were quantified by fluorometry using Qubit^®^ Protein Assay Kit. From those solutions, an equivalent aliquot of 50 μg protein was transferred to 0.5 mL tubes and dried down. The protein pellets were suspended in 6 M Urea aqueous solution (25 μL). The same volume of reducing reagent plus 10 mM DTT was added, and the samples were reduced for 60 min at room temperature. Subsequently, 50 μL of the alkylation solution, 100 mM IAA, was added, and the samples were alkylated for another 60 min at 54 °C in the dark. Afterward, 1M DTT (1 μL) was added to react with the remaining IAA. Finally, 100 μL ice-cold trypsin solution at 1:50 (trypsin/protein) ratio was added to the samples, followed by incubation for 16 h at 37 °C. Following digestion, the reaction was stopped by adding of 10% formic acid (5 μL). The samples were then desalted using ZipTips^®^ and kept at −20 °C until the analysis.

Peptides were analyzed by on-line nanoflow LC-MS on an EASY-nLC II system (Thermo Scientific) connected to an LTQ-Orbitrap Velos instrument (Thermo Scientific) via a Proxeon nanoelectrospray ion source. Peptides were separated on an analytical EASY-Column (10cm, ID75μm, 3μm, C18-Thermo Scientific) previously trapped in a pre-column EASY-Column (2cm, ID100μm, 5μm, C18-Thermo Scientific). Tryptic digested peptides were separated using a 60-min linear gradient of 0–60% buffer B (acetonitrile in 0.1% formic acid) at 300nL/min flowrate. The LTQ-Orbitrap Velos mass spectrometer was operated in positive ion mode using DDA (data-dependent acquisition) mode. Full MS scans were performed with 60,000 of resolution, and the m/z range for MS scans was 400–1200. The minimum signal threshold was 15000 counts and, for dynamic exclusion, it was considered as 1 repeat count with a duration of 30 s. To discriminate the charge state of the peptides, the charge state screening was enabled, and ions either with unassigned charge state or singly charged were rejected.

The MS/MS spectra from each LC-MS/MS run were searched against five different databases with two distinct search engines, an in-house Proteome Discoverer 1.4 software (Thermo, USA). The search criteria were as follows: full tryptic specificity was required, two missed cleavage was allowed, carbamidomethylation (C) was set as the fixed modification, and the oxidation (M) was set as the variable modification, precursor ion mass tolerances were set at 10 ppm for all MS acquired in an orbitrap mass analyzer, and the fragment ion mass tolerance was set at 0,6 Da for all MS2 spectra acquired. All covariates were log-transformed before statistical analysis. All the analyses were performed using STRING software and UniProt for protein-protein interactions, identification, and statistics. P ≤0.05 was considered significant.

### Oxygen consumption rates

An hour before oxygen consumption measurements, cell media was replaced by assay media (2 mM glucose, 0.8 mM Mg^2+^, 1.8 mM Ca^2+^, 143 mM NaCl, 5.4 mM KCl, 0.91 mM NaH_2_PO_4_, and 15 mg/mL Phenol red) for 60 min at 37°C (no CO_2_) before loading into the Seahorse Bioscience XF96 extracellular analyzer. During the 60 min period, the ports of the cartridge containing the oxygen probes were loaded with the compounds to be injected during the assay (75 μL/port), and the cartridge was calibrated. Basal respiration was recorded for 30 min, at 4 min intervals, until system stabilization. FCCP was used at final concentrations of 5 mM and injected with sodium pyruvate (Sigma) at a final concentration of 5 mM. Oligomycin and antimycin A were used at final concentrations of 1 and 10 μg/mL, respectively. Rotenone was used at a concentration of 1 μM. All respiratory modulators were used at ideal concentrations titrated during preliminary experiments (not shown). A typical OCR chart is displayed, where OCR represents the percentage of basal respiration.

### Western blotting

Proteins derived from cell lysates were separated on SDS-PAGE gel (4 to 12%) (Novex, Invitrogen) with 2-(N-morpholino) ethanesulfonic acid (MES) buffer (Novex, Invitrogen) at 150V for 60 to 90 minutes. Proteins were loaded onto Polyvinylidene difluoride (PVDF) membranes (Millipore, Temecula, CA, USA) in buffer containing 10% Tris-glycine and 15% methanol. The proteins were transferred at 32 V for 90 minutes after pre-treatment for 1-2 minutes with methanol. Membranes were blocked with 3% BSA containing Tris buffer in saline solution (TBS) for 60 minutes. After blocking, the membranes were incubated with specific primary antibodies in TBS containing 3% BSA and 0.1% Tween-20 overnight. Primary antibodies: goat anti-mouse Naip1 (sc-11067, 1:200, Santa Cruz Biotech., Santa Cruz, CA, USA), rabbit anti-mouse NLRP3 (mAb #15101, 1:1000, Cell Signalling Technology, Danvers, MA, USA), anti-β-actin mouse (1:1000, Li-COR Bioscience, Lincoln, NE, USA), and anti-IL1β from R&D ELISA kit, detection antibody (1:1000). Membranes were washed and incubated with secondary antibodies (IRDye 800CW or IrDye 680RD, LI-COR Biosciences, 1:20000) for 60 minutes. After washing, the staining was visualized on the Odyssey imaging system. The quantification of staining was performed using the Fiji / Image J program.

### Confocal imaging

Confocal Laser Scanning Microscopy (CLSM) was performed with a Leica TCS SP5 SMD confocal system (Leica Microsystems). The images were collected using a single z step, and the emitted fluorescence was detected by scanned detectors with 490–520-nm, 575– 605-nm, and emission filters. Pre-defined settings for laser power and detector gain were used for all experiments. Microphotographs were analyzed using the LAS AF version 2.2.1 (Leica Microsystems) or Volocity 6.01 software.

### Quartz Crystal Microbalance (QCM)

The interaction of the Naip1 protein with the sUA was analyzed in a QCM device (SRS, Stanford Research Systems). A plasmid carrying Naip1 GFP-tagged was virally transduced into THP-1 cells and cell lysates were used to bind the Naip1 protein into the anti-GFP, previously immobilized onto gold quartz crystals (SRS, 5 MHz). Prior to the antibody immobilization, gold crystals were immersed in piranha solution 1:3H2O2/H2SO4 for 15 min, washed twice with absolute ethanol for 5 min, followed by three times washing with ultrapure water for 5 min, and dried with a gentle flow of nitrogen. Afterward, gold crystals were incubated with EDC (100 mmol/L) and NHS (150 mmol/L) and then, 200 μL of anti-GFP (MA5-15256, Thermo Fisher Scientific), at 20 μg/mL, diluted in ultrapure water was deposited over each crystal for 16 h at 4 °C in a humid chamber. The crystals were rinsed in three sequential ultrapure water baths, dried at 22 °C, and blocked with 1% BSA solution for 1 h at 37 °C. After the incubation, all sample chips were washed and dried, as described above. Then, 200 μL of cell lysates were also deposited for 16 h at 4 °C in a humid chamber. The crystals were placed in a QCM flow chamber apparatus connected to a syringe pump with a 100 μL min-1 flow rate (KD Scientific). An initial hydration step with 500 μL of ultrapure water was done, and 500 μL of each sUA solution sample (12.5 μM; 25 μM; 50 μM; 100 μM; and 200 μM) was injected in individual experiments. Each experiment was done in triplicates, and the results were expressed as the average value.

### Surface plasmon resonance (SPR)-based immunosensor development

An Autolab Sprit instrument (Eco Chemie B. V., The Netherlands), which presents the phenomenon of attenuated total internal reflection (Kretschmann configuration) as operation mode [73] was employed in the SPR analysis. This SPR technique is equipped with a glass prism (BK7), a planar gold SPR sensor chip, and two measurement channels (channel 1 and channel 2). For the measurements, a laser diode with a wavelength fixed at 670 nm and a photodiode detector were employed. In terms of functionality, changes near the interface metal/environment promote a change in the resonance conditions of the system as a result, a shift in the θ_SPR_ occurs. In this sense, SPR techniques allow obtaining information on biomolecular interactions in real time.

The experiments were performed as demonstrated by Souto *et* al. [74]. Previously to the gold surface functionalization, the SPR sensor chip was cleaned in piranha solution (1:3 mixture of 30 % H_2_O_2_ and concentrated H_2_SO_4_) for approximately 1 minute, followed by the immersion of the substrate in acetone (5 minutes), and then in isopropyl alcohol (5 minutes). After this, the SPR sensor chip was washed with deionized water and dried with a pure N_2(g)_ flow. The functionalization of the gold surface was performed through the formation of a self-assemble monolayer (SAM), which was obtained by the reaction for 24 hours of ethanolic solution consisting of 11-mercaptoundecanoic acid (11-MUA, 1.0 mmol L^−1^). After the formation of the film on the gold surface (11-MUA/Au), it was copiously washed with ethanol, water, and dried with N_2(g)_ flow. All steps described above were performed *ex situ*. In the next step, the functionalized SPR sensor chip was immediately inserted into the SPR instrument, and the measurements were performed in real time. The terminal carboxyl groups of 11-MUA were activated via PBS buffer solution (10 mmol L^−1^ at pH 7.4) consisting of EDC (100 mmol/L) and NHS (150 mmol/L) for approximately 10 minutes for the formation of the NHS-ester groups. This strategy was used to allow the covalent binding of anti-GFP (20 μg/mL) on the SAM onto gold (11-MUA/Au). Afterward, successive additions of the buffer solution were performed to remove the excess of molecules onto the surface. Then, the immobilization of the anti-GFP was monitored via the SPR technique for approximately 45 minutes. This step was accompanied by washed step by using the successive addition of buffer solution. To prevent non-specific binding, after the immobilization of the anti-GFP on the 11-MUA/Au (SAM/Au), the unbound reactive ester groups were deactivated by the PBS buffer solution consisting of ethanolamine - EA (1.0 mol L^−1^ at pH 8.5) for approximately 5 minutes. Successive additions of the buffer solution then removed the excess of unbounded EA molecules. After successfully characterizing the immobilization and blocking steps, the interaction between the anti-GFP and Naip1 protein (GFP-tagged) was evaluated. Cell lysate derived from GFP-tagged THP-1 cells expressing mNaip1 was used to immobilize Naip1. After characterizing the interaction between anti-GFP and Naip1 protein, an aqueous solution of sUA was added to evaluate its interaction with Naip1 protein. As control assay, the interaction of palmitate with Naip1 protein was also evaluated.

### Protein structure analysis

The homology models were obtained using MODELLER v.9.18 [75] with the structure 4KXF as a template [76].To fix residues with bad torsion angles, the target proteins were repaired using RepairPDB [77], and Chimera [78] was used to add hydrogen atoms and charges where adequate. Surface electrostatic potentials were calculated using the AMBER force field implemented on APBS [79], taking as input files converted on PDB2PQR [80]. For each NAIP structure, a blind docking was performed on SwissDock [81], having the uric acid on its ionized form (Urate) as a ligand (ZINC AC: 2041003). During the docking, the surfaces of both proteins were scanned for putative binding pockets in more than 250 iterations. Several low energy ligand clusters with similar binding modes (poses) were found. All poses showing ΔG < −6 kcal/mol were considered in further analyses, and those showing the lowest energies were selected as the best representation of the binding between urate and both human and murine Naip.

### Statistics

Experiments were performed in duplicate or triplicate and at least two independent tests were performed for each assay. The data were described in terms of the mean and S.E.M. unless specified in the figure legend. Differences between groups were compared using ANOVA (with Tukey’s post-test) and Student’s *t*-test. Significant differences were regarded as p<0.05, p<0.01 or p<0.001, according to the figure. All statistical analyses were performed using GraphPad PRISM 6.01 (La Jolla, CA, USA).

## Acknowledgements

We thank Prof. Dario Simoes Zamboni for providing us femurs and tibias from WT, Naip1^−/−^, Naip2^−/−^, Naip5^−/−^, ΔNaip^−/−^ and Nlrc4^−/−^ mice. We also thank Uira Souto Melo for all discussion concerning the data present in the paper.

---

**Supplemental Figure 01.**
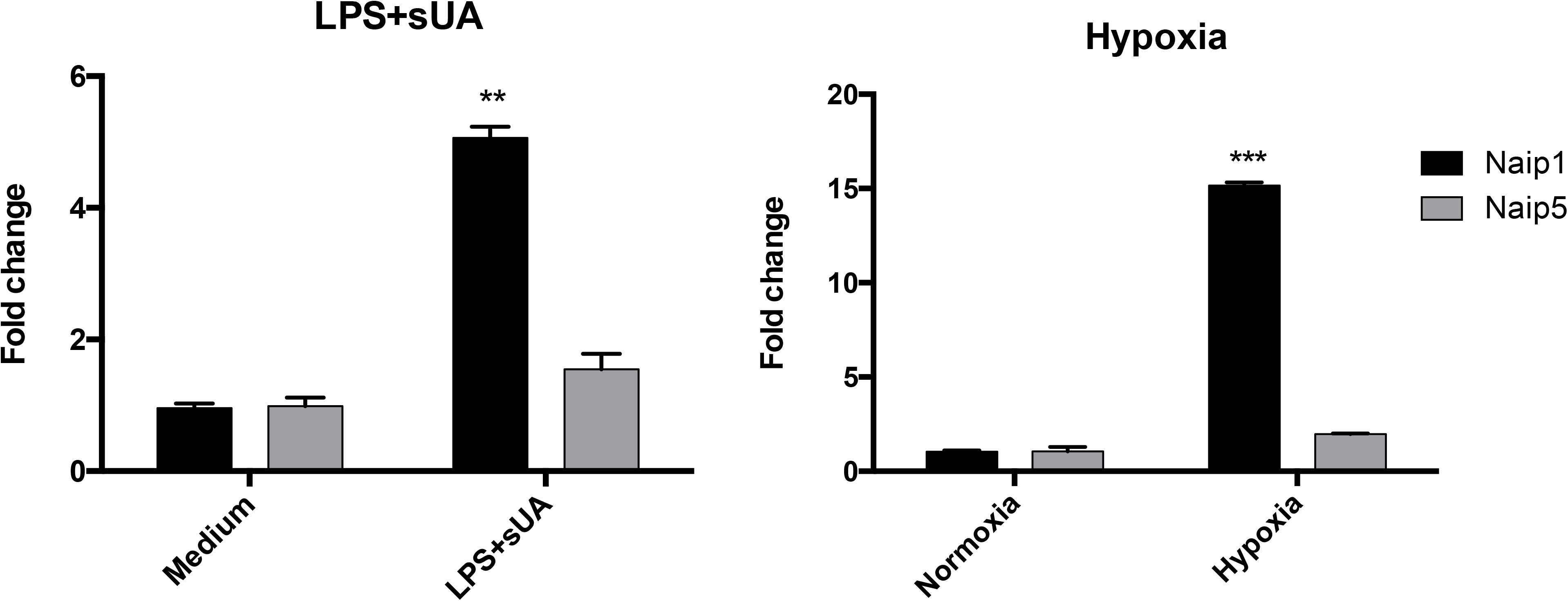
Soluble uric acid induces Naip1, but not Naip5, mRNA expression. Naip1(black bars) and Naip5 (gray bars) mRNA levels in bone marrow-derived macrophages after (**A**) sUA stimulus for 24 hours into LPS-primed cells when compared to non-treated cells, and (**B**) hypoxia condition for 24 hours, when compared to normoxia condition. qPCR data were normalized to HPRT expression, and the mean expression in (A) non-treated cells and (B) normoxia condition was considered 1. n = 3 animals per group in each experiment. **p < 0.01, ***p < 0.001.

**Supplemental Figure 02.**
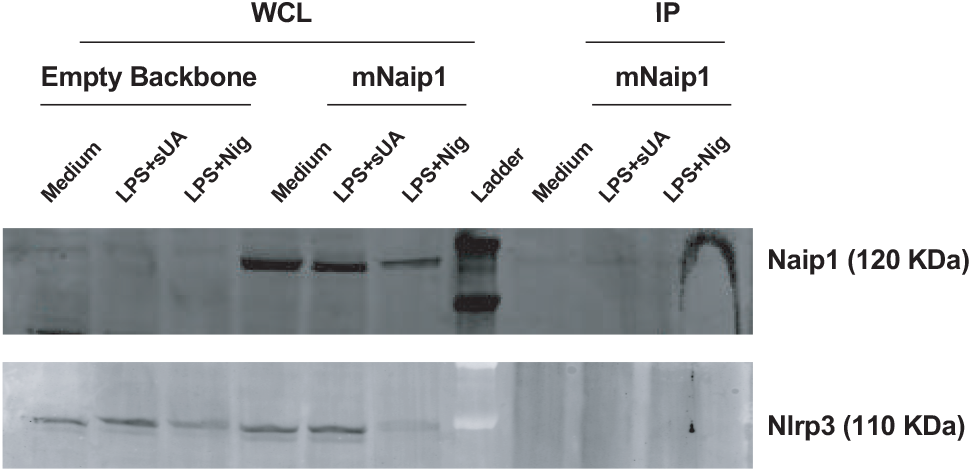
Nlrp3 and Naip1 may not directly interact to each other upon sUA stimulus. Immunoprecipitation - western blot analysis on THP1 whole cell lysates (WCL) and on THP1 immunoprecipitants (IP) using Nlrp3 mAb (upper panel) and Naip1 polyclonal Ab (lower panel) followed by anti-Naip1 and anti-Nlrp3 antibodies, respectively. Cells were analyzed under three different conditions: non-stimulated (medium), LPS-primed and sUA-stimulated (LPS+UA), and LPS-primed and nigericin-stimulated (LPS+Nig). Figures are representative of three different experiments.

**Supplemental Figure 03.**
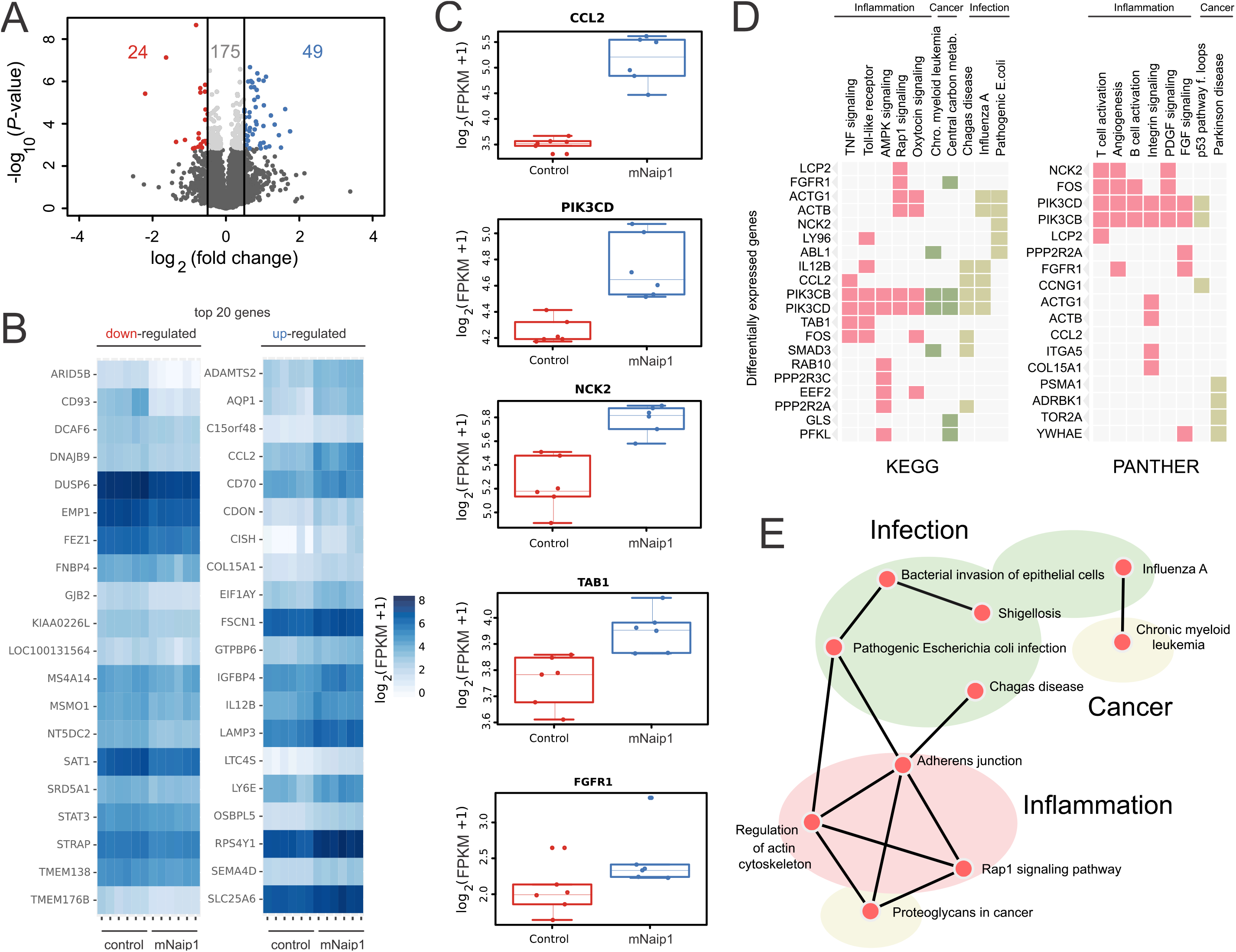
THP1 cells transduced with mNaip1 shows increased expression of inflammation-related genes after sUA treatment. THP1 cells were transduced with a lentiviral vector containing NAIP1 and GFP (NAIP1) and only with GFP gene (control). Cells were sorted on the basis of their GFP fluorescence followed by LPS pretreatment and sUA treatment then mRNA was purified and sequenced. (**A**) Volcano plot of gene expression changes. The y-axis specifies the negative logarithm to base 10 of t-test *P*-values and x-axis specifies the logarithm of fold changes to base 2. Colored dots indicate significantly differential expressed genes (q-value < 0.05). Vertical lines reflect threshold criteria for up-regulated genes (log2 fold change > +0.5) colored in blue (n=49), down-regulated genes (log2 fold change < −0.5) colored in red (n=24), and not differentially expressed genes in light grey (n=175). (**B**) Top 20 most down and up-regulated genes in NAIP1 cells (q-value < 0.05) by means of centered logarithm of FPKM values in the six replicates of each experimental group. (**C**) Box-plot showing FPKM expression levels (q-value < 0.05) of selected genes *CCL2* (NM_002982), *PIK3CD* (NM_005026), *NCK2* (NM_003581), *TAB1* (NM_006116), and *FGFR1* (NM_023106) in control and NAIP-1 cells. (**D**) Heatmap of enriched KEGG 2016 and Panther 2016 database terms as columns and differentially expressed (q-value < 0.05 and n=248) genes as rows. Enriched terms (*P*-value < 0.05) were grouped as inflammation, infection or cancer-related major groups. (**E**) Network of enriched terms on KEGG database found in enrichment analysis of only up-regulated genes (q-value < 0.05, fold change > 1, and n=136). Each node represents a KEEG 2016 term and links represents that a term (node) have some genes in common. KEEG terms are highlighted in inflammation, infection and cancer major groups. The experiments were performed in duplicates in three independent analysis.

**Supplemental Figure 04.**
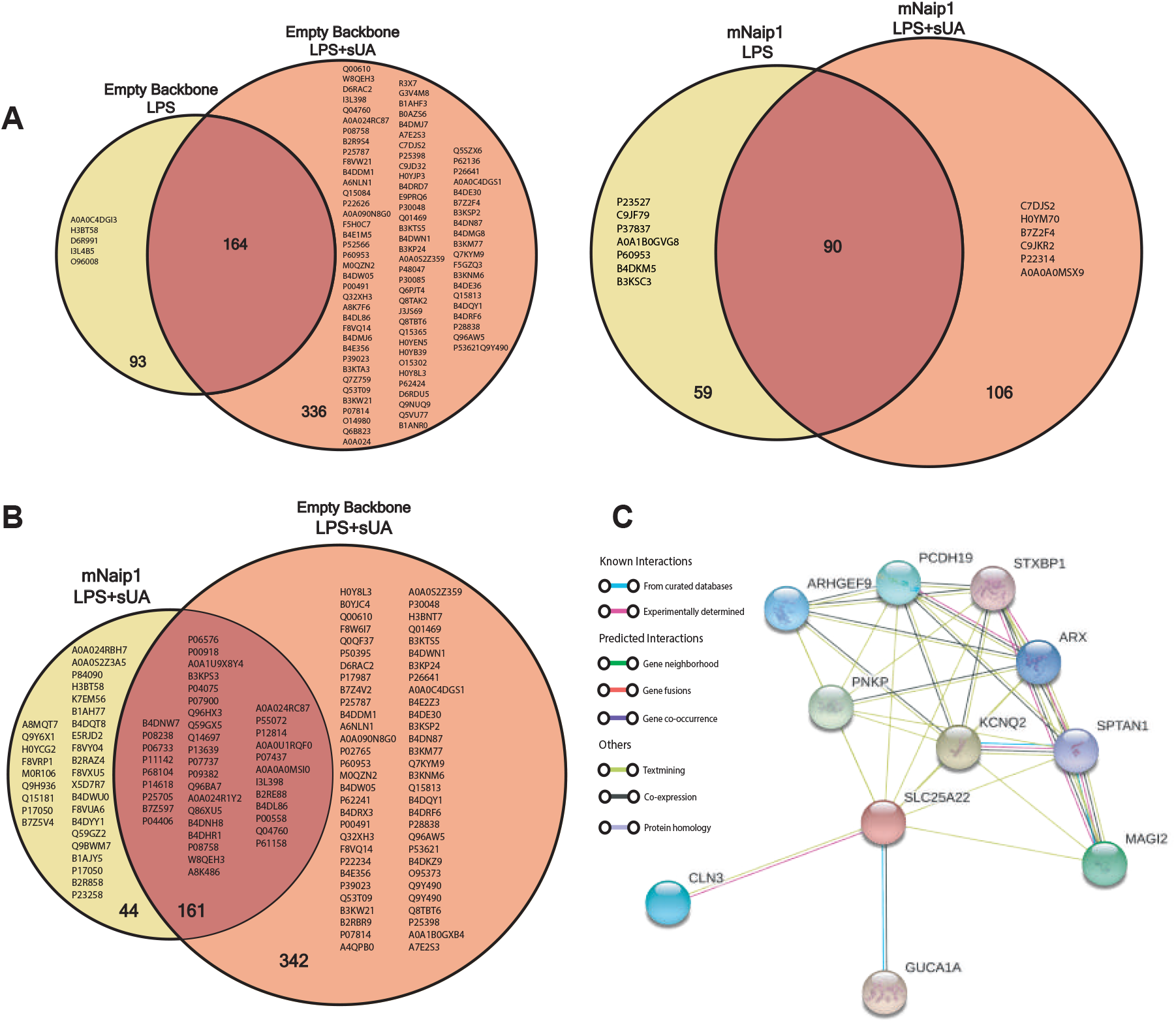
sUA triggers altered cellular protein content in cells expressing mNaip1. (**A**) Proteomic analysis of LPS-primed or LPS-primed and sUA-stimulated macrophages transduced with empty backbone (left panel) or mNaip1 (right panel) showing proteins only present in each condition. The numbers indicate the amount of proteins exclusively expressed in each condition. (**B**) Proteomic analysis of LPS-primed and sUA-stimulated macrophages showing thirty proteins only present in mNaip1-expressed cells (in the yellow circle) and sixty proteins only present in empty backbone-transduced cells (in the orange circle). The numbers indicate the amount of proteins exclusively expressed in each condition. (**C**) STRING network view of proteomics analysis showing upregulated proteins in mNaip1-expressing cells versus empty backbone-transduced THP-1 ones, both after sUA stimulus. Colored lines between the proteins indicate the various type of interaction evidence. The experiments were performed in triplicates.

**Supplemental Figure 05.**
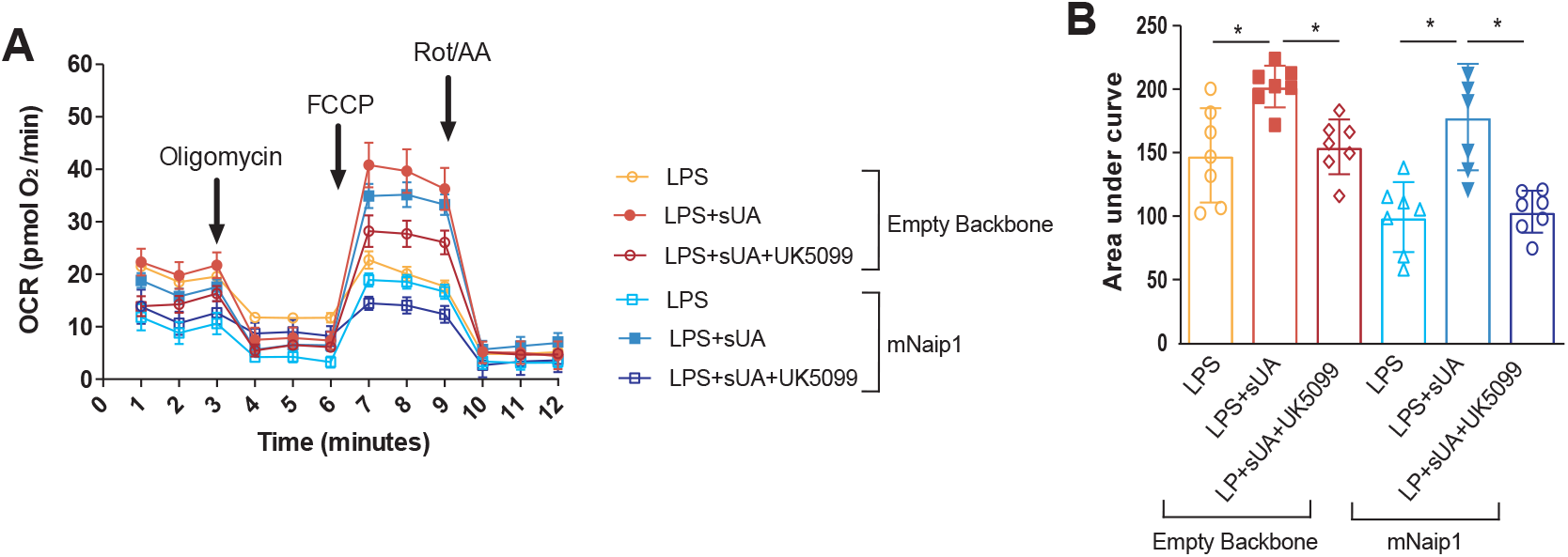
sUA triggers increased maximal respiration in a mNaip1-independet way. (**A**) Bioenergetic profiles of THP-1 cells under different stimuli. Cells (60,000 per well) were treated with respiratory inhibitors and uncoupler at the following concentrations: oligomycin (1 μg/mL), CCCP (5 μM) and antimycin A (10 μg/mL) plus rotenone (1 μM). The graph shows representative oxygen consumption rates (OCR) from LPS-primed control cells (light red line), LPS-primed and sUA-stimulated control cells (red line), LPS-primed and sUA-stimulated control cells under UK5099 (100 μM) pre-treatment (dark red line), LPS-primed Naip1 expressing cells (light blue line), LPS-primed and sUA-stimulated Naip1 expressing cells (blue line), and LPS-primed and sUA-stimulated Naip1 expressing cells under UK5099 pre-treatment (purple line). (**B**) Area under curve of graph in A. Data are representative of three independent experiments and n = 7 for each analyzed condition. *p < 0.05.

